# Cortical Tracking of Speech and Music Predicts Reading Ability in Adults

**DOI:** 10.64898/2026.02.18.706526

**Authors:** Sarah C. Allen, Stratos Koukouvinis, Saara M. Varjopuro, Anne Keitel

**Affiliations:** Psychology, University of Dundee, Dundee, Scotland; Department of Psychology, University of Turku, Turku, Finland; School of Psychology and Neuroscience, University of Glasgow, Glasgow, Scotland

## Abstract

Cortical tracking of acoustic features is essential for the neural processing of continuous stimuli such as speech and music. For example, it has been shown that children with dyslexia show atypical cortical tracking. This tracking may therefore reflect a fundamental auditory temporal processing mechanism supporting literacy more generally. In the current pre-registered study, we tested the hypothesis that cortical tracking of speech and music predicts reading ability in healthy young adults (N = 32), evaluated through a lexical decision task. Participants first completed an online session in which they performed a lexical decision task to assess their reading skills. This was followed by an electroencephalography (EEG) session, in which participants listened to a naturalistic short story and a music track. Using mutual information, we showed that neural activity aligned to both speech and music across a wide range of frequencies. Interestingly, cortical tracking was stronger for speech at very low frequencies, while it was stronger for music at higher frequencies. Critically, cortical tracking predicted reaction times in the lexical decision task in a frequency-dependent manner: stronger delta-band tracking (∼1-3 Hz) for both speech and music was associated with faster reaction times, whereas stronger alpha-band tracking (∼12 Hz) for speech was associated with slower reaction times. These findings remained significant even when controlling for stimulus type, age, musical experience and reading enjoyment. These results suggest that cortical tracking of speech and music reflect a domain-general temporal processing mechanism that is associated with reading ability beyond stimulus-specific features, and beyond development. These findings advance the neurobiological underpinnings of literacy and could potentially be leveraged for developing new reading interventions.

## INTRODUCTION

Speech and music are (quasi-) rhythmic signals formed of hierarchically organised structural units that unfold over time (Lerdahl & Jackendoff, 1983; Miyagawa et al., 2013; Asano, 2022). For speech, these units include phonemes and syllables, while for music, they include notes and motifs. Aside from their rhythmic structure, speech and music share other characteristics: they are both auditory signals structured by acoustic cues, such as duration (timing), frequency (pitch) and amplitude (loudness).

Changes in these acoustic cues are reflected by the modulation of the signal’s amplitude over time (Patel, 2010), often quantified as the acoustic envelope (Peelle & Davis, 2012). Neural activity aligns with these slow envelope fluctuations of continuous external stimuli, known as cortical tracking (Giraud & Poeppel, 2012; Doelling & Poeppel, 2015; Ding et al., 2017). Whether this alignment is due to synchronisation of endogenous oscillations to an external rhythm or to evoked responses is still under debate (Haegens & Golumbic, 2018; Obleser & Kayser, 2019), see Atanasova et al. (2026) for a comprehensive review. Some factors which influence cortical tracking are attention (Symons et al., 2021), language and musical proficiency (Doelling & Poeppel, 2015), stimulus familiarity (Kumagai et al., 2017; Keitel et al., 2025), and prior knowledge (Sohoglu et al., 2012).

Neural activity in the brain tracks the envelopes of speech and music at multiple timescales (Giraud & Poeppel, 2012; Doelling & Poeppel, 2015; Ding et al., 2017). In speech, different temporal frequencies have been associated with various linguistic units: syllables are largely associated with theta (4-8Hz), and words and phrases with delta (0.5-4Hz) (Ahissar et al., 2001; Peelle & Davis, 2012; Gross et al., 2013; Keitel et al., 2018). A right hemisphere bias for processing slow (syllabic/prosodic) information has been found (Abrams et al., 2008), whereas the left hemisphere preferentially extracts information from faster temporal features (Obleser et al., 2008), consistent with the asymmetric sampling hypothesis (Poeppel, 2003). Cortical tracking has been linked to language and literacy development, though the precise nature of this relationship remains complex. Research with dyslexic populations has yielded varied findings, with some studies reporting reduced tracking of low-frequency acoustic information (corresponding to syllabic and prosodic rhythms) and others demonstrating enhanced tracking at higher frequencies (Lehongre et al., 2013; Molinaro et al., 2016; Power et al., 2016). These frequency-specific results may have direct implications for reading development, as reading relies upon cognitive mechanisms that parallel those underlying speech perception. Specifically, graphemes (clusters of letters), must be mapped onto phonological representations to enable fluent decoding (Goswami, 2007). The temporal sampling framework (Goswami, 2011) proposes that reading difficulties in dyslexia stem from impaired cortical tracking of speech at multiple temporal scales. It is unknown whether cortical tracking predicts reading ability in healthy individuals.

Music, like speech, contains hierarchical temporal structures such as rhythm and meter, which are also cortically tracked (Doelling & Poeppel, 2015; Harding et al., 2019; Keitel et al., 2025). These cortical tracking mechanisms are thought to overlap with those observed in speech processing, particularly those used for parsing syllables and prosodic patterns, suggesting a shared temporal processing system (Harding et al., 2019), although it is currently still under discussion to what extent speech and music processing are domain-specific or domain-general (Kotz et al., 2018; Drakoulaki et al., 2024).

Due to rhythm deficits often found in dyslexic populations, researchers have investigated the role of music (experience and training) related to language and reading, with many studies finding that musical training leads to improved reading skills in children (Tierney & Kraus, 2013; Garcia-de-Soria et al., 2025). In particular, rhythm reproduction has been found to be strongly associated with phonological awareness (Bhide et al., 2013; Goswami et al., 2013; Flaugnacco et al., 2014), however, causality remains contested. A recent review proposed that observed differences in perceptual and cognitive abilities between musicians and non-musicians reflect pre-existing aptitude differences (leading to selection bias) rather than effects of far transfer (Schellenberg & Lima, 2024).

Here, we investigated whether cortical tracking of continuous natural speech and music, quantified through mutual information, is associated with reading skills in healthy adults, quantified through a lexical decision task, while controlling for age, musical sophistication and reading enjoyment. We expected that stronger cortical tracking of the speech and music envelopes would predict better performance in the lexical decision task (see preregistration: https://osf.io/t4dje/).

## MATERIALS AND METHODS

### Participants

Thirty-two volunteers participated in this study (17 female, 19-28 years old; *M* = 22.2, *SD* = 2.50). Participants declared never having received a diagnosis of neurological/psychological disorders or dyslexia. All participants were right-handed (Oldfield, 1971). Self-reports of hearing ability (Five-minute Hearing test, revised version) (Koike et al., 1994) indicated that 31 participants had no hearing impairments, while one participant was recommended a hearing test (score of 23/60, with 20 being the cutoff). This study was approved by the School of Social Sciences Research Ethics Committee at the University of Dundee (approval number: UoD-SoSS-PSY-UG-2021-263) and adhered to the guidelines for the treatment of human participants in the Declaration of Helsinki. Volunteers received monetary compensation of £10/h. Research questions, hypotheses, measured variables and analyses were pre-registered on the OSF website (https://osf.io/xrq36). Deviations from the pre-registration are detailed where necessary.

### Procedure

#### Online procedure

Prior to taking part in the EEG experiment, participants were asked to complete the first part of the study online, using the experiment builder Gorilla (Anwyl-Irvine et al., 2020). In this online session, participants completed a demographics questionnaire, a musicality assessment (Müllensiefen et al., 2013), a handedness questionnaire (Oldfield, 1971) and the lexical decision task. The musicality questionnaire included 14 items from the Goldsmiths Musical Sophistication Index (GMSI) covering perceptual abilities, active engagement, singing abilities and musical training. Each item included a statement such as “I spend a lot of my free time doing music-related activities”, which participants rated on a scale of 1 to 7. The online task and questionnaires had to be completed on a PC and in a single, uninterrupted session.

#### In-person procedure

After completing the online portion of the study, participants were invited to take part in the in-person EEG session. Participants performed the EEG experiment in a soundproof booth (160 cm x 110 cm) where they were seated approximately 65cm away from a 24-inch monitor. Participants were equipped with high-quality, wired headphones (Sennheiser, HD 25, 75Ω). The experiment was run using Psychtoolbox (version ‘3.0.17)(Brainard & Vision, 1997; Pelli & Vision, 1997) through MATLAB (MATLAB, 2021). During the presentation of speech and music stimuli, participants were instructed to relax, limit motor movement to avoid movement artefacts and fixate on a small green circle on the screen. Blocks were self-paced, and the order of the auditory stimuli was counterbalanced.

### Stimuli and tasks

#### Lexical Decision Task

The lexical decision task was adapted from a pre-existing task (no reference was found for the task or the study in which it was used) on the experiment-building website Gorilla (Anwyl-Irvine et al., 2020). The words and non-words used in this task were taken from a previous study (Yao et al., 2018) and are part of the corpus of Glasgow Norms (Scott et al., 2019). Both words and nonwords ranged from 3 to 11 letters in length (*M* = 6.11, *SD* = 1.75). All words were concrete nouns with neutral valence, as words with differing emotional connotations can affect response times (Yao et al., 2018). Word frequency (occurrences per million as per the British National Corpus) was on average *M* = 23.95 (*SD* = 36.49). In the task, participants were asked to use the left and right arrow keys to indicate whether the string of letters on the screen was a word (e.g., ‘statue’) or a non-word (e.g., ‘depane’, which does not have any meaning in the English language). Participants were given 12 practice trials with feedback. At the beginning of each block, there were two brief screens, appearing for 1000 ms each, saying “Ready?” and “GO!” to prepare the participant for the trials. A fixation cross was also shown for 500 milliseconds before each word and nonword in the task. The experimental task consisted of 90 trials, given in 2 blocks of 45 trials each, and feedback was not provided. All tasks and questionnaires used Open Sans font. Halfway through the lexical decision task, participants were given the option to take a break and recommence when they preferred.

#### Passive Listening to speech and music

All auditory stimuli were presented at a sampling rate of 44,100 Hz. The short story selected for this study was “The Elves and the Shoemaker”, originally written by the brothers Grimm, read by a female speaker with a pleasant voice (https://librivox.org/). The length of the story was 300 s (5 minutes). The articulation rate of the stimulus was calculated using Praat (Boersma, 2001), through the automatic detection of syllable nuclei based on intensity peaks in the speech signal (De Jong & Wempe, 2009). The stimulus had an articulation rate of 3.61 syllables per second. Similarly, the modulation spectrum of the speech stimulus showed a peak at 3.75 Hz (Figure 1A).

**Figure 1.**
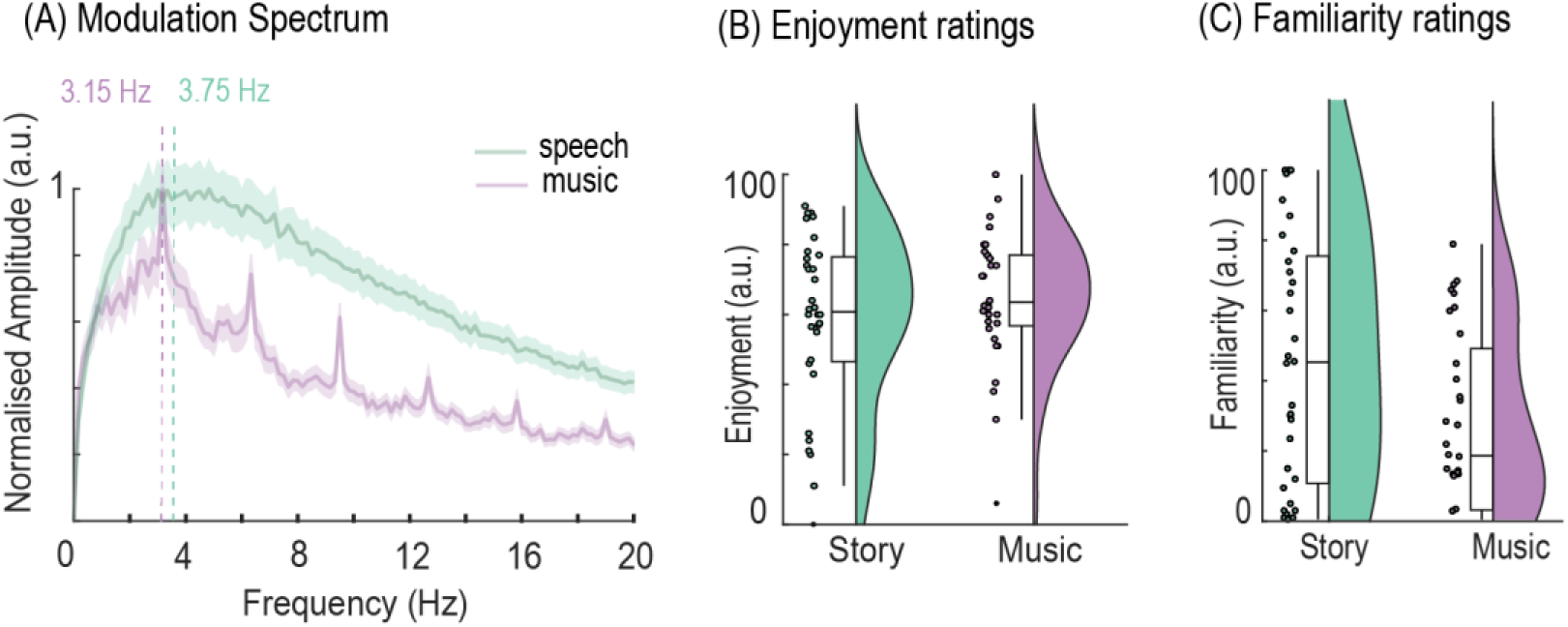
(A) Modulation spectrum of speech (green) and music (purple) stimuli. Thick lines indicate average values across 6-s segments, and shaded areas represent the standard error of the mean. Peak frequency is shown with dotted lines. (B) Enjoyment ratings for speech and music excerpts. There was no significant difference between stimulus ratings (*p* = .287). (C) Familiarity ratings for speech and music excerpts. The story was rated as more familiar than the music piece (*t*(31) = 2.55, *p* = .015). *Note*: Dots in (B) and (C) show individual data points, violin plots show kernel density estimates, and boxplots show median interquartile ranges and minimum/maximum.

The music piece selected was “Fluid” by Lin Rountree, a pleasant jazz piece at 95 BPM. This piece featured bass, keyboard and trumpet; there were no vocals included. The duration was 255 s (4 minutes and 25 seconds), and the modulation spectrum showed a peak at 3.15Hz (Figure 1A). After each piece, participants rated how familiar and how pleasant they had found the stimuli by using a Visual Analog Scale: this involved placing a vertical marker between “did not enjoy at all” and “enjoyed a lot” for enjoyment and “not at all familiar” and “extremely familiar” for familiarity (each analysed in arbitrary units between 0 and 100). Participants rated the story (*M* = 45.67, *SD* = 35.17) as significantly more familiar than the music piece (*M* = 27.27, *SD* = 25.17) (*t*(31) = 2.55, *p* = .015, Figure 1C). However, enjoyment ratings did not differ significantly between the story (*M* = 58.72, *SD* = 24.38) and the music piece (*M* = 64.80, *SD* = 18.98) (*t*(31) = -1.08, *p* = .287, Figure 1B). All stimuli are available on the OSF server (https://osf.io/xrq36).

### Analysis of behavioural data

For analysis of the lexical decision task, incorrect trials (*M* = 5.69, *SD* = 3.25) were excluded from analysis. For correct trials, responses faster than 250 ms or exceeding 1500 ms were excluded (*M* = 6.58, *SD* = 10.45), as these could reflect inadvertent movements or lapses in attention (Yao et al., 2018). Three participants were excluded from further analyses involving the lexical decision task as their total number of rejected trials exceeded 2 *SD*s from the group mean. For the remaining participants, median reaction times and accuracy rates were calculated separately for word and nonword trials. These values were then averaged to yield total accuracy and reaction time measures.

### Acoustic envelope pre-processing

To analyse the neural tracking of speech and music signals, the wideband envelope of the stimuli was extracted. The acoustic waveforms were filtered into eight frequency bands (between 100 and 8000 Hz, 3rd order Butterworth filter, forward and reverse) equally distant on the cochlear frequency map (Smith et al., 2002). The signal in each of these frequency bands was then Hilbert-transformed and the magnitude extracted before being averaged to obtain the wideband music and speech envelopes used in further analyses. Lastly, envelopes were down-sampled to a sampling rate of 150 Hz (Keitel et al., 2018).

### EEG acquisition and pre-processing

EEG was recorded from 64 scalp electrodes and digitally sampled at 512 Hz, using a BioSemi ActiveTwo system. Scalp electrodes were positioned according to the international 10-20 system. Electrodes with an offset of greater/less than ±20 mV were adjusted prior to starting the recording. Ultimately, electrode offset for all electrodes was below an absolute value of 30 mV before the experiment began. Lateral eye movements were monitored by two electro-oculographic electrodes placed at the outer canthus of each eye. Vertical eye movements and blinks were monitored by two electro-oculographic electrodes positioned below and above the left eye.

Data pre-processing was conducted using the Fieldtrip Toolbox (Oostenveld et al., 2011) and custom-made scripts in MATLAB (R2025a) (The Mathworks, 2025). The data were cut according to the length of the stimuli, with an additional 2-s leading and trailing window. Data was first re-referenced to Cz, then, fourth-order Butterworth low-pass (60 Hz) and high-pass (0.2 Hz) filters were applied. Noisy (e.g., with higher variance than most) channels were visually identified and interpolated through triangulation. A maximum of five channels was interpolated per participant (*M* = 1.25, *SD* = 1.34). To remove eye artifacts and blinks, an independent component analysis (ICA) was then carried out for 30 principal components. Components, including eye movements or blinks, were selected and removed (*M* = 1.40, *SD* = 0.49). The EEG signal was then down-sampled to 150 Hz to match the envelope signals (Keitel et al., 2018).

### MI analysis

The correspondence between the continuous EEG signal and the acoustic envelope signals was analysed using a Gaussian copula mutual information framework (Ince et al., 2017). The Mutual Information (MI) between the continuous L1-normalised EEG signals and the envelopes of the auditory stimuli (as well as their derivatives, see Fig. 2) was calculated in the frequency domain by applying a continuous wavelet transform (Chalas et al., 2022), for 63 logarithmically spaced frequencies between .25 and 20 Hz. We used a participant-specific optimal brain-stimulus lag computed by identifying, for each participant and condition, the lag at which the MI value was highest (identified at electrode *Cz* for slow frequencies, averaged between 0.5 – 4 Hz). Each MI value was computed per participant, condition, frequency and channel. To normalise MI, we first created random (surrogate) MI distributions. For this, we segmented the continuous envelope signals into 5-s segments and shuffled the segments randomly. This kept the statistical properties of the signal but destroyed the temporal relationship between the acoustic stimuli and brain signals. MI was then computed between the brain signal and the shuffled envelope signals. Normalised MI was computed by z-scoring the observed MI values against a surrogate distribution, subtracting the mean and dividing by the standard deviation of randomised MI estimates. The normalised MI values were used for all further analyses.

**Figure 2.**
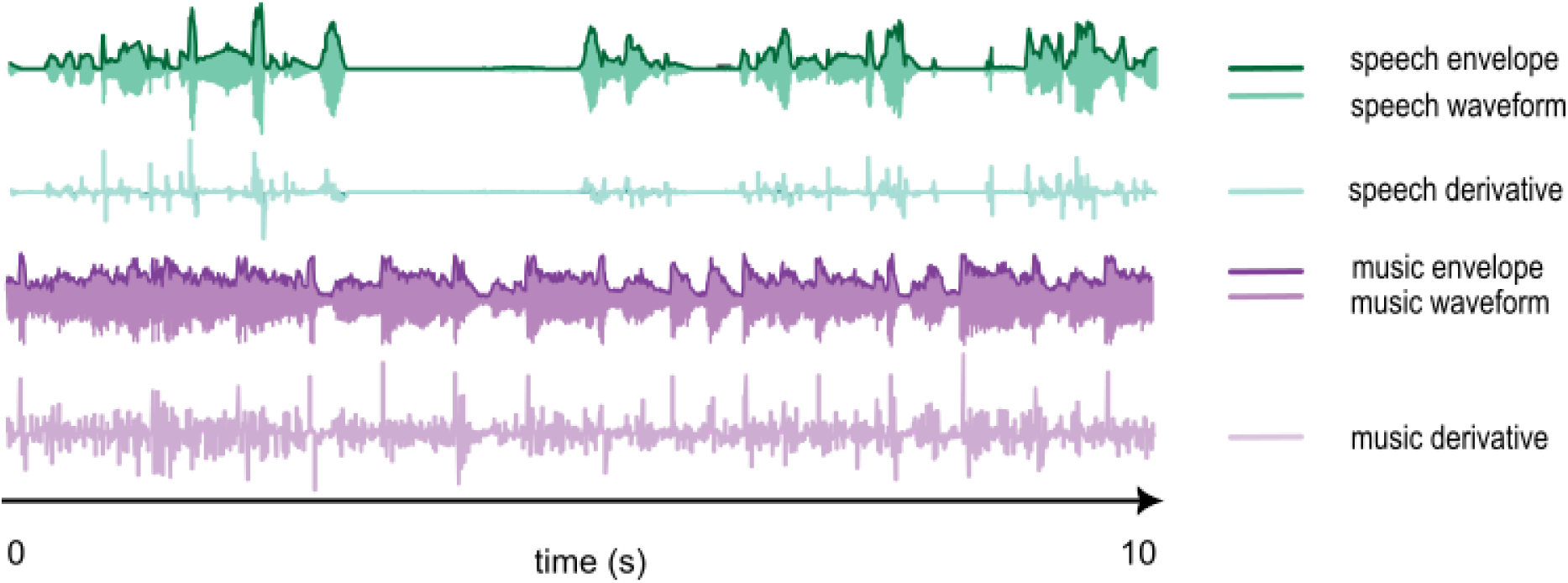
Brief excerpt (10 s) of acoustic envelope, waveform, and first derivative for the speech (green) and music stimuli (purple).

### Statistical analysis

For each participant, normalised MI values at each channel and frequency were tested against a null baseline of zero using dependent-samples t-tests. Cluster-based multiple-comparison correction (implemented in FieldTrip (Oostenveld et al., 2011)) was applied using 5000 Monte Carlo permutations, in which the condition labels (normalised MI vs. zero) were randomly exchanged within subjects to form the null distribution. Clusters were defined as spanning more than 3 frequency bins, and the cluster-level statistic was the sum of *t*-values. Observed clusters were considered significant if their cluster statistic exceeded the 95th percentile of the permutation distribution. This controlled for the family-wise error rate at the cluster level. To avoid reporting spurious results, we only report clusters that cover a range of frequencies of more than 0.5 Hz.

For the comparison between the two tracking conditions, dependent samples *t*-tests were computed across channels and frequencies with MI values of one condition compared against MI values of the other, with the same cluster-permutation approach used for multiple comparison as described above.

To test the relationship between cortical tracking and behavioural measures (reaction times from the lexical decision task), Pearson’s correlations were computed between the behavioural measures and the normalised MI values, across all electrodes and frequencies. Before comparing the *r* values with the permutation distribution using cluster-based permutation (using a minimum cluster size of 3 channels and tested against the 95^th^ percentile of the permutation distribution), Pearson’s *r* values were transformed to be normally distributed using Fisher’s z-transformation (Gorsuch & Lehmann, 2010). As an indicator of effect sizes, we report Cohen’s *d* for peak electrodes.

To further compare the contribution of cortical tracking in both speech and music conditions on performance in the lexical decision task, a robust multiple linear regression was computed (R 4.5.1). The model included: (i) MI values, (ii) stimulus type (speech or music), (iii) frequency band (delta or alpha), (iv) musical sophistication scores, (v) age, and (vi) reading enjoyment as predictors, along with the three-way interaction between MI × stimulus type × musical sophistication and the two-way interaction between MI × frequency band. Reaction times in the lexical decision task served as the outcome variable. All continuous variables were z-scored. Additionally, we performed a Bayesian model comparison (Wagenmakers, 2007) to evaluate the evidence for or against the presence of an interaction effect (MI × stimulus type). Specifically, we computed Bayesian linear regression models that only differed in whether the interaction was included.

To explicitly test for hemispheric lateralisation in our significant correlation clusters, we computed a lateralisation index (LI) following the procedure outlined by Haegens et al. (2011) which is comparable to the method used by Thut et al. (2006). The LI was calculated as:

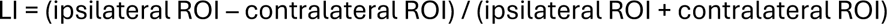

where the region of interest (ROI) comprised the electrodes within each significant lateral cluster. For each participant, correlation values (*r*) from electrodes in one hemisphere were paired with their hemispheric counterparts. The resulting LI values were then tested against zero using a two-sided one-sample *t*-test to determine whether correlations differed significantly between hemispheres.

## RESULTS

### Behavioural results

In the word recognition task, a paired *t*-test found that reaction times to words (*M* = 658.53 ms, *SD* = 75.68) were significantly faster than to nonwords (*M* = 777.13 ms, *SD* = 106.87), *t*(28) = -8.68, *p* < .001 (see Figure 3A). Accuracy was high for both words (*M* = 94.38%, *SD* = 3.83) and nonwords (*M* = 92.87%, *SD* = 6.56) with no significant difference between conditions, *t*(28) = 1.04, *p* = 0.304 (Figure 3B). The preregistration included both accuracy and reaction times as indicators of reading ability. However, given that accuracy was at ceiling level and there was no significant difference between conditions, we used only reaction times for the following analyses.

**Figure 3.**
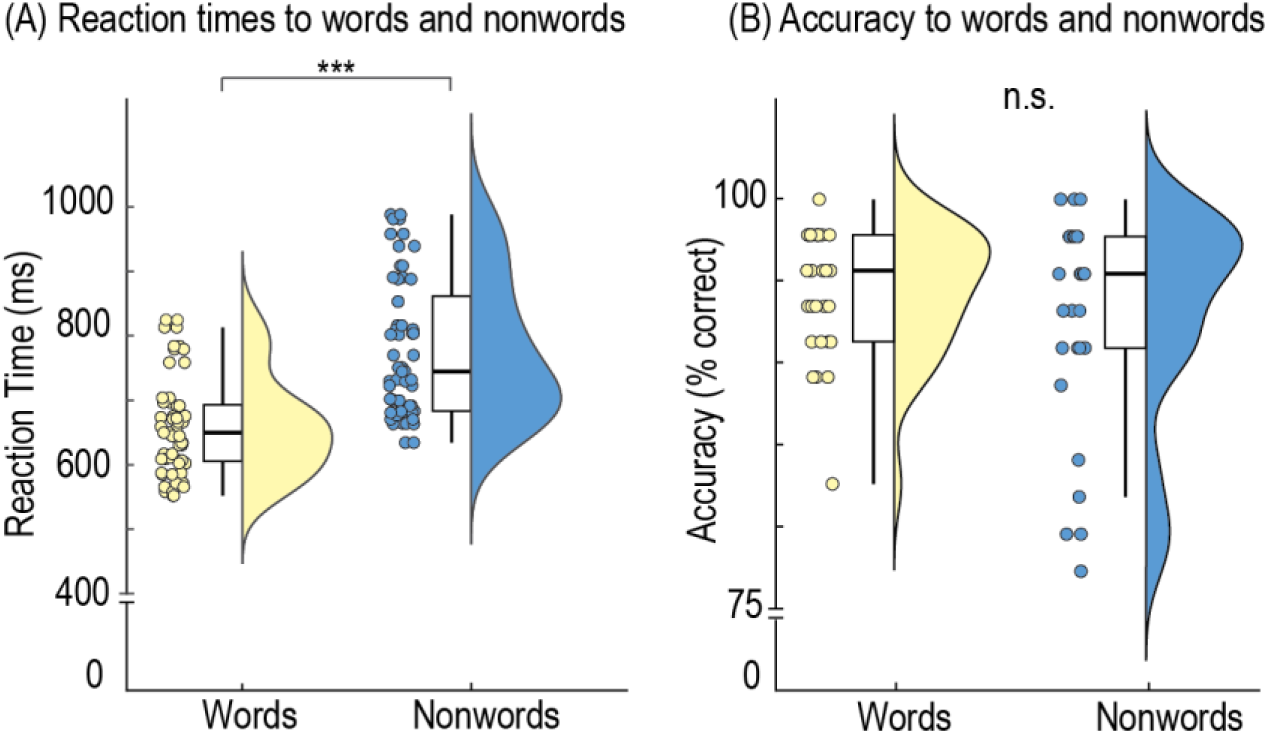
Violin plots and boxplots showing median reaction times to words and nonwords (A) and accuracy for words and nonwords (B). Points indicate individual data for all participants; violin plots show kernel density estimates and boxplots show median interquartile ranges and minimum/maximum.

### Cortical tracking of the speech and music envelopes

We analysed whether participants tracked the acoustic speech and music envelopes using dependent-samples *t*-tests of the normalised MI values (across channels and frequencies) against a baseline of zero, with cluster-based permutation to control for multiple comparisons. For speech (Figure 4A), we found a large positive cluster of all 64 electrodes, ranging from 0.26 Hz to 12.37 Hz, which significantly tracked amplitude fluctuations with a peak at 0.67 Hz (Cohen’s *d*_peak_ = 2.59, *p*_cluster_ < .001). Similarly, for music (Figure 4B), we found a large positive cluster of all 64 electrodes that significantly tracked amplitude fluctuations from 0.26 Hz to 18.75Hz with a peak at 5.38Hz (Cohen’s *d*_peak_ = 3.43, *p*_cluster_ < .001). This indicates widespread spatial and spectral tracking of both speech and music stimuli.

**Figure 4.**
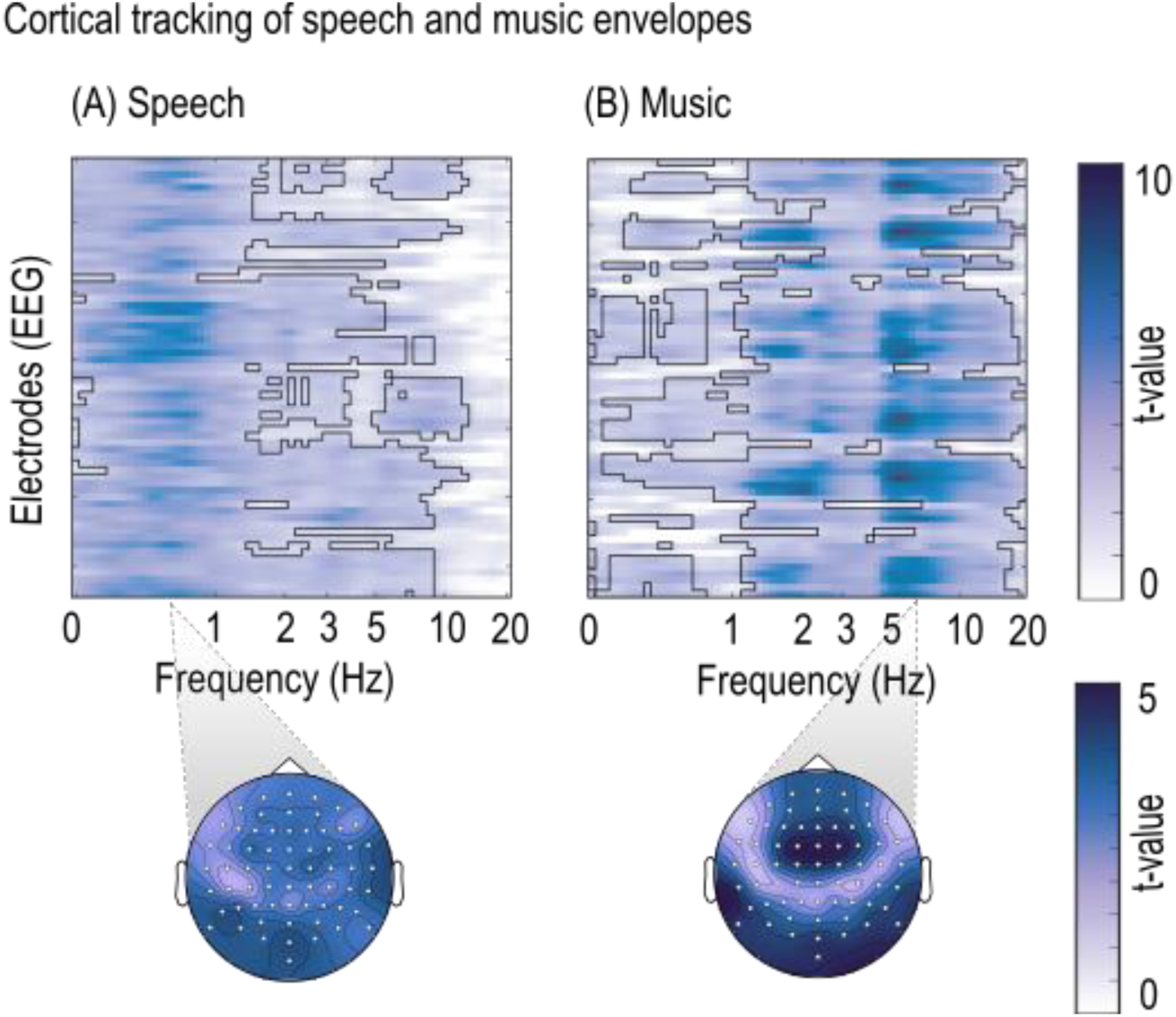
Cortical tracking of acoustic speech (A) and music (B) envelopes. Channel-by-frequency heatmaps of *t*-values indicating cortical tracking, with significant clusters outlined in black. Below each heatmap is the topography of the average cortical tracking across all significant frequencies with significant channels highlighted in white.

### Comparison of speech and music tracking

We then directly compared cortical tracking in speech and music conditions across channels and frequencies using dependent-samples *t*-tests with cluster-based permutation to control for multiple comparisons. Three significant clusters emerged: one negative cluster and two positive clusters (Figure 5). A large negative cluster of 56 electrodes (Figure 5B(i)), ranging from 0.26 Hz to 1.09 Hz (Cohen’s *d*_peak_ = 1.87, *p*_cluster_ < .001) indicated that the speech envelope was tracked more strongly than the music envelope at low frequencies. The two positive clusters indicate that the music envelope was tracked more strongly than the speech envelope. The first positive cluster (Figure 5B(ii)) included 18 temporal and occipital electrodes and ranged from 2.34 Hz to 3.55 Hz (Cohen’s *d*_peak_ = 1.48, *p*_cluster_ = 0.032) while the second positive cluster (Figure 5B(iii)) included 59 electrodes and ranged from 1.66 Hz to 16.32 Hz (Cohen’s *d*_peak_ = 2.82, *p*_cluster_ < .001).

**Figure 5.**
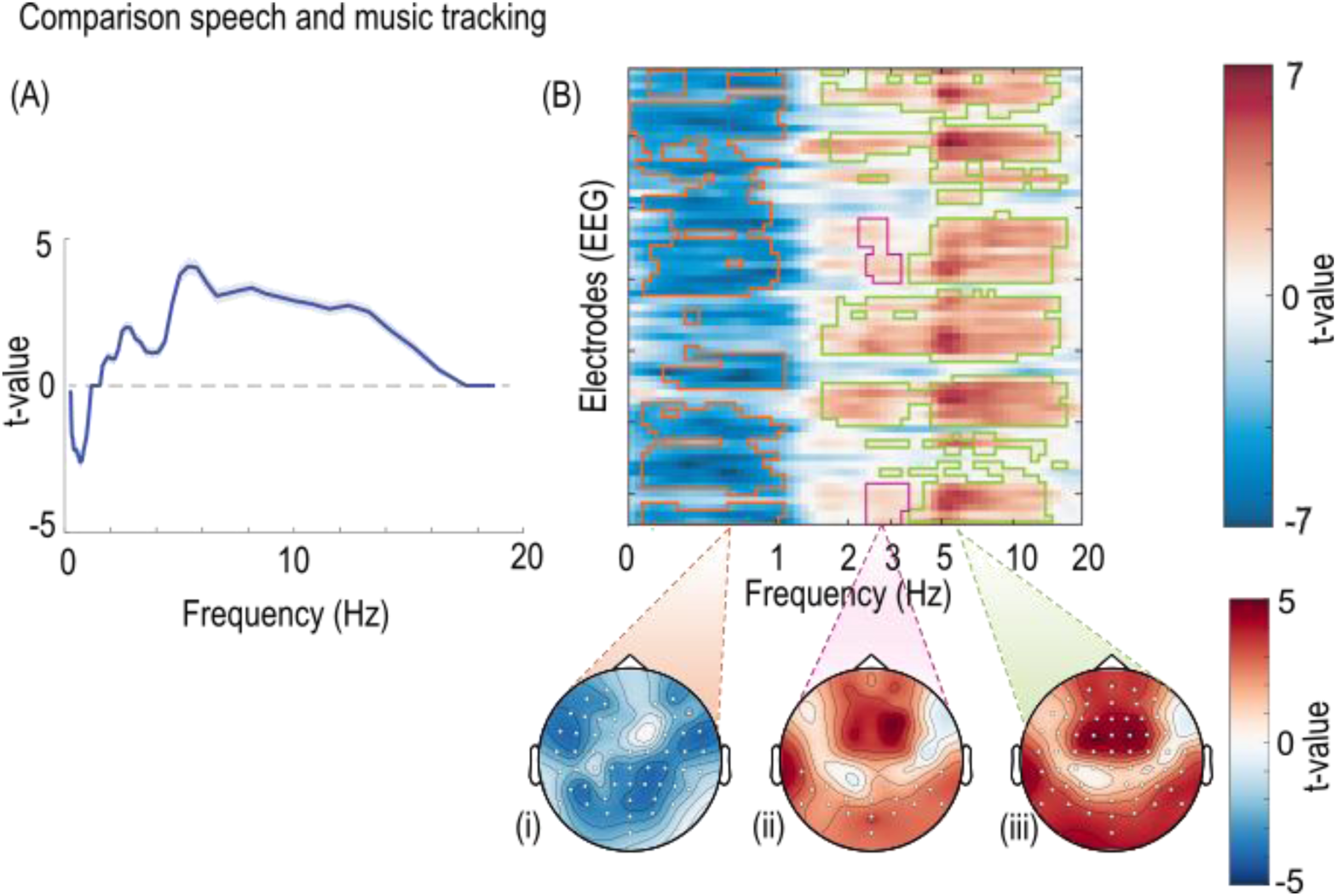
(A) Mean *t*-values for the comparison between cortical tracking of speech and music across all frequencies. Negative values indicate stronger tracking of speech (at low frequencies between 0.26 and 1.09 Hz). Positive values indicate stronger tracking for music (at frequencies between 1.66 and 16.32 Hz). Shaded areas indicate the standard error of the mean. (B) Channel-by-frequency heatmap of *t*-values of the comparison between cortical tracking of speech and cortical tracking of music. The negative cluster (i), showing stronger cortical tracking of speech, is shown in orange. Positive clusters, showing stronger tracking of music, are shown in pink (ii) and green (iii). Topographies show the average cortical tracking across all significant frequencies, with significant channels shown in white.

### Cortical tracking predicts performance in the lexical decision task

To test whether envelope tracking during listening to the story and music predicted participants’ behavioural performance in the lexical decision task, we correlated the MI values per electrode and frequency with participants’ average reaction time across words and nonwords. Note that separate initial analyses for reaction times for words and nonwords indicated that results were comparable (Figure S1); therefore, we opted to use median reaction times across all trials to streamline the presentation of results.

For the story condition, we found one negative occipital cluster (Figure 6A(i)) at low frequencies (0.89 Hz - 1.78 Hz), that significantly predicted reaction times in the lexical decision task (Cohen’s *d*_peak_ = 1.39, *p*_cluster_ < .001, 9 electrodes), which was right lateralised *t*(11) = -3.09, *p* = .010, indicating that participants who showed stronger envelope tracking to speech at low frequencies had faster reaction times in the lexical decision task.

**Figure 6.**
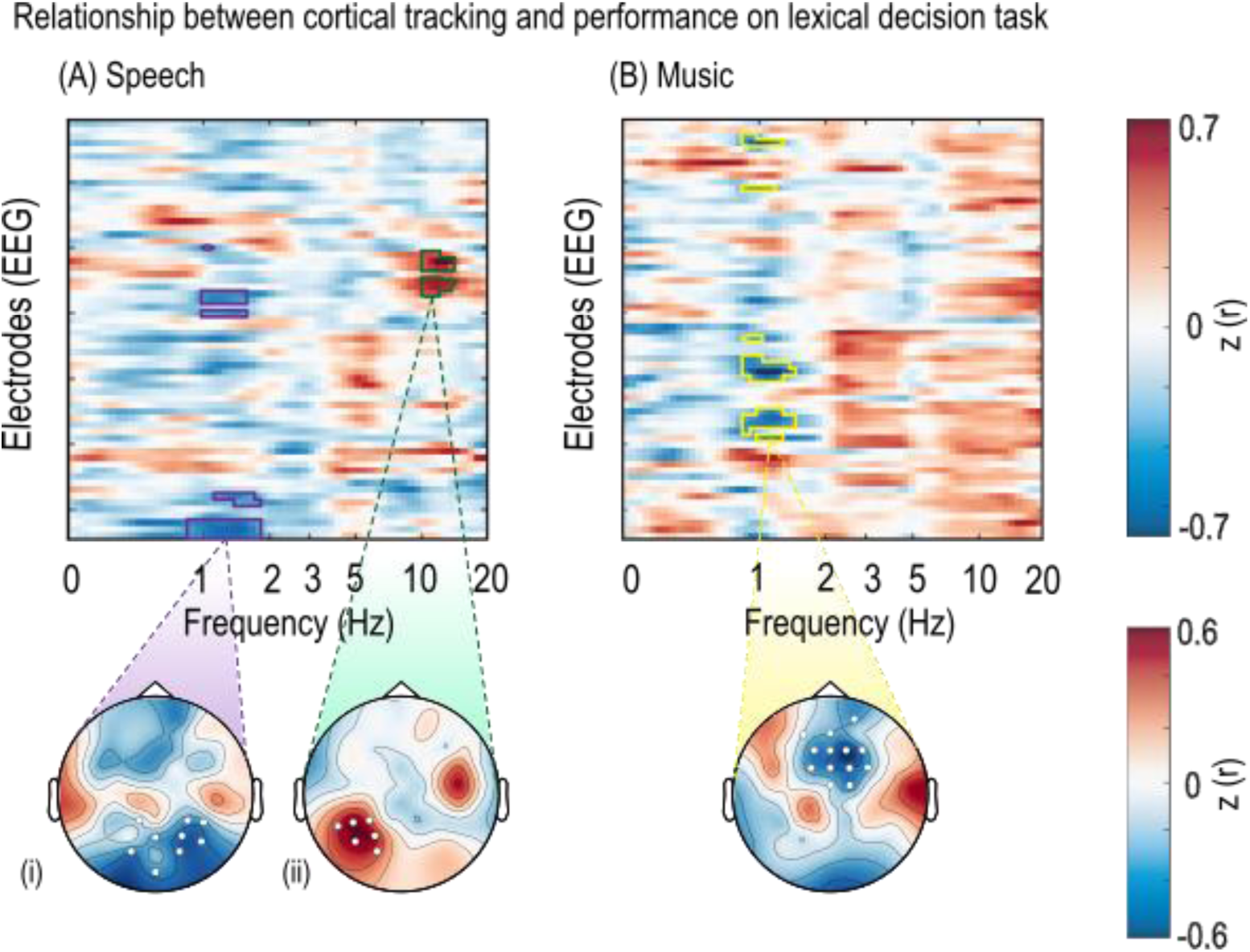
Relationship between cortical tracking and performance on the lexical decision task, quantified through reaction times. (A) Channel-by-frequency heatmap of the correlation between cortical tracking of speech and participants’ response speed in the lexical decision task (reaction times). Two clusters were found for speech: each topography shows the average *r*-values across frequencies of the cluster. The negative cluster (i) indicated that stronger tracking of speech correlated with faster reaction times in the lexical decision task. The positive cluster in the left hemisphere in occipital electrodes at high frequencies (ii) indicated that stronger tracking of speech correlated with slower reaction times in the lexical decision task. (B) Channel-by-frequency heatmap of the correlation between cortical tracking of music and participants’ response speed in the lexical decision task. The topography below shows the average *r*-values across frequencies of the negative cluster. Here, stronger tracking of music was associated with better performance (i.e., faster reaction times). *Note*: Significant channels are highlighted with white circles.

We also found one positive cluster (Figure 6A(ii)) in occipital and parietal electrodes at high frequencies (10.05 Hz – 13.26 Hz) that predicted reaction times in the lexical decision task (Cohen’s *d*_peak_ = 2.41, *p*_cluster_ < .001, 6 electrodes), which was left lateralised (*t*(25) = 9.155, *p* < .001). This indicates that participants who showed stronger envelope tracking to speech at those higher frequencies had slower reaction times in the lexical decision task.

For the music condition, we found one negative cluster in frontal electrodes at low frequencies (0.89 Hz – 1.44 Hz) that predicted reaction times in the lexical decision task (Cohen’s *d*_peak_ = 2.40, *p*_cluster_ < .001, 13 electrodes; Figure 6B). This indicates that participants who showed stronger envelope tracking to music at low frequencies had faster reaction times in the lexical decision task.

To assess the joint contribution of cortical tracking in both speech and music conditions to lexical decision performance, and control for additional variables such as musical sophistication, reading enjoyment and age, we performed an additional multiple linear regression in R (Version 4.5.1). The model included: (i) MI values averaged within positive/negative significant clusters, (ii) stimulus type (speech or music), (iii) frequency band (delta or alpha), (iv) musical sophistication scores, (v) age, and (vi) reading enjoyment as predictors, along with the three-way interaction between *MI × stimulus type × musical sophistication* and the two-way interaction between *MI × frequency*. Reaction times in the lexical decision task served as the outcome variable (Table 1). Importantly, if speech and music tracking predicted reaction times differentially, we would expect a significant interaction between *MI × stimulus type*. The main effect of cortical tracking (Figure 7A) was statistically significant (*β* = - 0.46, 95% CI [-0.75, -0.17], *p* = .003), indicating that, overall, participants with stronger cortical tracking exhibited faster reaction times in the lexical decision task. This confirms the results of the main analysis (Figure 6), while controlling for participant-related variables. Neither the main effect of stimulus type (*p* = .687) nor musical sophistication (*p* = .299) reached significance, suggesting that these factors did not directly influence reaction times.

**Figure 7.**
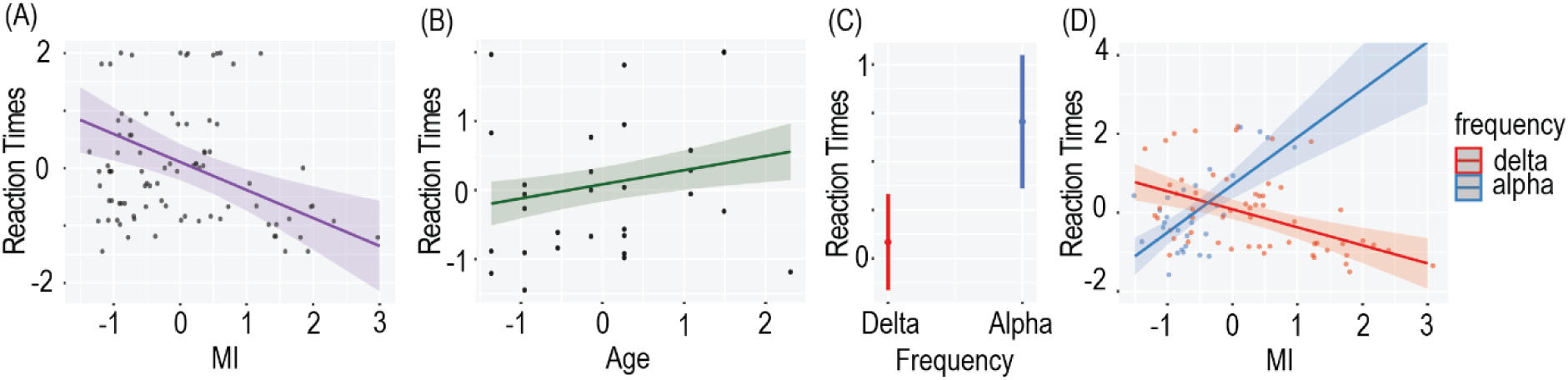
Relationship between cortical tracking and performance on the lexical decision task. (A) Main effect of MI on reaction times, with the purple line indicating the prediction model and the dots showing individual data points. (B) Main effect of age on reaction times, with the green line indicating the prediction model and the dots showing individual data points. (C) Main effect of frequency band (alpha vs delta) on reaction time. Dots indicate estimated marginal means from the regression model, and whiskers show 95% confidence intervals. (D) Two-way interaction: participants with higher MI in the delta frequency band also show faster reaction times, whereas participants with higher MI in the alpha frequency band show slower reaction times in the lexical decision task.

**Table 1.** Summary table for the robust linear regression model predicting reaction times.

| Predictors | Estimates | SE | 95% CI | | $p$ |
| --- | --- | --- | --- | --- | --- |
|  |  |  | LL | UL |  |
| Intercept | .08 | .16 | -.25 | .41 | .614 |
| **MI** | -.46 | .14 | -.75 | -.17 | **.003 \*\*** |
| Stimulus [Music] | -.10 | .24 | -.56 | .37 | .687 |
| Musical Sophistication | -.13 | .12 | -.36 | .11 | .299 |
| **Frequency [alpha]** | .62 | .28 | .06 | 1.18 | **.031 \*** |
| **Age** | .20 | .10 | .02 | .39 | **.034 \*** |
| Reading Enjoyment | .05 | .32 | -.14 | .24 | .623 |
| MI x Stimulus [music] | .11 | .23 | -.34 | .56 | .616 |
| MI x Musical Sophistication | .01 | .13 | -.24 | .26 | .910 |
| Stimulus [music] x Musical Sophistication | -.15 | .21 | -.58 | .27 | .473 |
| **MI x Frequency [alpha]** | 1.67 | .32 | 1.02 | 2.32 | **<.001 \*\*\*** |
| MI x Stimulus [music] x Musical Sophistication | .23 | .21 | -.19 | .65 | .280 |

Note. Significant predictors are highlighted in bold. \* $p < .05$ , \*\* $p < .01$ , \*\*\* $p < .001$

Age showed a significant main effect (Figure 7B; *β* = 0.20, 95% CI [0.02, 0.39], *p* = .034), with older participants showing slower reaction times. Frequency also showed a significant main effect (Figure 7C; *β* = 0.62, 95% CI [0.06, 1.18], *p* = .031) with cortical tracking in the alpha band showing slower reaction times than in the delta band. The interaction between cortical tracking and frequency (Figure 7D) was statistically significant (*β* = 1.67, 95% CI [1.02, 2.31], *p* <.001), indicating that the relationship between MI and reaction times was dependent on the frequency band. Specifically, stronger cortical tracking in the delta band was associated with faster reaction times, whereas stronger tracking in the alpha band was associated with slower reaction times, as shown in the main analysis (Figure 6) while controlling for musical sophistication, age and reading enjoyment. No other two-way or three-way interactions were statistically significant (all *p-*values > .280). Notably, the interaction between MI and stimulus did not reach significance, which might suggest that speech and music tracking did not contribute differently to reading skills. To formally evaluate evidence for or against the *MI × stimulus* interaction, we conducted a Bayesian model comparison (Wagenmakers, 2007). Specifically, the original model including the interaction was compared to a model that was identical, except for not including the *MI × stimulus* interaction. The Bayes factor indicated that the data were 5 times more likely under the model without interaction than under the original model (BF₁₀ = 0.19), providing substantial evidence against the interaction.

## DISCUSSION

The present study examined the relationship between cortical tracking of continuous naturalistic speech and music and reading skills, quantified through a lexical decision task. Several key findings emerged. First, cortical tracking predicted reading performance in a frequency-dependent manner: stronger delta-band tracking was associated with faster reaction times, whereas stronger alpha-band tracking was associated with slower reaction times. These findings remained even when controlling for stimulus type, musical sophistication, age and reading enjoyment. Secondly, both speech and music were found to be extensively tracked across a wide range of channels and frequencies. Finally, cortical tracking of speech was found to be stronger than that of music at low frequencies (<1 Hz), while cortical tracking of music was found to be stronger at higher frequencies (1-16Hz).

### Domain-general cortical tracking predicts reading performance

Cortical tracking of speech has been identified as a potentially critical mechanism underlying language and literacy development, as suggested by studies demonstrating atypical tracking patterns in children with dyslexia (Power et al., 2016; Araújo et al., 2024). We here tested whether cortical tracking of speech, as well as music, would predict individual differences in reading skill in healthy adults. Consistent with this hypothesis, cortical tracking emerged as a significant predictor of reading performance.

There was no main effect of stimulus type on lexical decision task performance, suggesting that both speech and music tracking contributed similarly to performance on the lexical decision task. Furthermore, Bayesian modelling provided substantial evidence in favour of the model without interaction. Taken together, our findings suggest that domain-general auditory processing mechanisms, rather than stimulus-specific features, underlie the relationship between cortical tracking and reading proficiency. Notably, the topographies and frequency ranges (Figure 6A(i) and Figure 6B) look remarkably similar, although for speech the effect reaches significance for posterior channels, while it reaches significance for music in anterior channels. The absence of a stimulus effect aligns with theoretical accounts proposing that shared temporal processing mechanisms support both linguistic and musical perception (Patel, 2011), and suggests that auditory processing mechanisms contribute to reading proficiency (Cason et al., 2015). We used a lexical decision task as a proxy for readings skill, as this has been shown to be strongly associated with reading proficiency (Weems & Zaidel, 2004; Lubineau et al., 2024). However, reading is a multi-faceted process, and future research should test whether the relationship between cortical tracking and reading skill also applies to other tasks, such as reading comprehension, reading fluency, and phonological decoding.

The relationship between cortical tracking and reading performance was modulated by frequency band. Stronger tracking in the delta band predicted faster reaction times, whereas stronger tracking in the alpha band predicted slower reaction times. This frequency-dependent pattern suggests that successful reading may depend on the integrated processing of multiple temporal scales. Enhanced delta-band tracking may facilitate the extraction of syllabic and prosodic information critical for phonological processing (Goswami, 2011) while excessive alpha-band tracking may reflect inefficient allocation of neural resources to fine-grained temporal details at the expense of syllabic-level information (Peelle & Davis, 2012). This interpretation is consistent with the temporal sampling framework, which posits that reading difficulties arise from impaired multi-scale temporal processing of speech (Power et al., 2016). The observed association between stronger alpha-band tracking and slower reaction times aligns with emerging evidence linking stronger high-frequency cortical tracking to reading difficulties. Individuals with dyslexia and developmental language disorder exhibit atypical frequency-specific tracking patterns (Araújo et al., 2024; Nora et al., 2024). Findings vary across studies: some report only reduced low-frequency tracking (Molinaro et al., 2016; Di Liberto et al., 2018) while others demonstrate enhanced high-frequency tracking (Lehongre et al., 2011). Given that slow and fast amplitude modulations are hierarchically nested in speech (Gross et al., 2013), atypical low-frequency temporal sampling could cascade to altered processing at faster rates (Keshavarzi et al., 2025). According to this, impaired extraction of syllabic-level information may require compensatory reliance on fine-grained temporal details (captured in alpha frequencies in our stimulus material), resulting in less efficient reading.

In the current study, we cannot make causal inferences about whether enhanced cortical tracking facilitates reading or whether better reading strengthens cortical tracking mechanisms. Additional research is needed to determine whether training-induced changes in cortical tracking would correspond to improvements in reading performance, or whether improvements in reading would correspond to changed cortical tracking, which would provide evidence for causal mechanisms. Additionally, reading proficiency comprises multiple component skills that may exhibit distinct relationships with cortical tracking. Future investigations should examine associations with specific subcomponents, including phonological decoding, reading fluency, prosodic sensitivity, and reading comprehension. This approach would clarify which specific aspects of reading ability are most closely linked to cortical tracking mechanisms.

### Lateralised relationship between cortical tracking of speech and reading proficiency

The pattern of results found for the predictive role of speech tracking, whereby the correlation of high-frequency cortical tracking with reading reaction times was left-lateralised, and the correlation of low-frequency cortical tracking with reading was right-lateralised (Figure 6A), is reminiscent of the Asymmetric Sampling in Time (AST) theory (Poeppel, 2003). According to the AST, the left hemisphere is proposed to be optimised for processing faster, fine-grained temporal aspects of speech, whereas the right hemisphere is proposed to be optimised for processing slower temporal features which unfold over time. While we did not find stronger speech tracking of high- and low frequencies in left and right hemispheres, respectively (consistent with previous studies, tracking was bilateral for both speech and music (Harding et al., 2019), Figure 4), the left- and right-lateralised *relationships* with reading performance might indicate that the processing of slower timescales in the right, and faster timescales in the left hemisphere, is relevant for higher-level cognition such as reading.

### Cortical tracking of speech is stronger than music at low frequencies

Previous work has shown that speech and music are tracked by cortical activity across multiple timescales (Giraud & Poeppel, 2012; Doelling & Poeppel, 2015; Ding et al., 2017). Therefore, we expected cortical tracking across a wide range of channels and frequencies. Our results found significant cortical tracking for both speech and music across a wide spatial distribution of channels and a broad spectral range. This extensive tracking pattern is consistent with previous research (Te Rietmolen et al., 2024) and is thought to reflect the hierarchical temporal structure of both domains, with different frequency bands capturing distinct structural units, including prosody and rhythm.

Research has found mixed results on the strength of speech versus music tracking. Some studies have shown that speech tracking is stronger than music tracking at low frequencies, corresponding to syllables and phrases (der Nederlanden et al., 2020), while others find speech tracking to be stronger at higher frequencies (Osorio & Assaneo, 2025). These discrepancies may stem partly from methodological differences in how frequency ranges are analysed. Unlike previous studies that averaged cortical tracking across frequency ranges (Harding et al., 2019), we used a frequency-resolved mutual information approach, enabling a more fine-grained comparison. This approach revealed that the relative strength of the cortical tracking is frequency-dependent: speech tracking was stronger than music tracking for very low frequencies (< 1 Hz), while music tracking was stronger than speech tracking across delta, theta and alpha bands. The stronger speech tracking at very low frequencies may reflect the tracking of prosodic phrases, which occur at rates below 1 Hz in natural speech. However, these frequency-specific differences could also be influenced by the particular acoustic characteristics of our stimuli, including differences in temporal regularity, rhythmic structure, and spectral content between the selected speech and music.

### Musical sophistication and reading enjoyment do not predict reading proficiency

Musical expertise has been associated with improved reading skills in children (Tierney & Kraus, 2013; Garcia-de-Soria et al., 2025), thought to be due to the positive effects of musical expertise on brain plasticity and development (Olszewska et al., 2021). However, evidence for this relationship in adults is mixed. While some studies report positive associations between musical training and phonological processing in adults (Pantaleo et al., 2024), others find no association between musical expertise and reading-related skills when controlling for general cognitive abilities (Swaminathan et al., 2018). Our results found that self-reported musical sophistication did not significantly influence performance on the lexical decision task. This suggests that potential transfer effects of musical training on language processing may be most pronounced during development. The GMSI (Müllensiefen et al., 2013) provides a broad assessment of musical sophistication, including self-reported engagement, training duration, and perceptual abilities. However, it should be noted that more specialised measures targeting specific perceptual skills such as pitch or rhythm discrimination may capture distinct dimensions of musical ability not fully reflected in self-report indices, potentially explaining differences in how musical experience relates to cortical tracking across studies. Furthermore, observed differences between musicians and non-musicians may not reflect the effects of musical training itself, but rather stem from pre-existing aptitude differences that influence who chooses to pursue musical training, leading to selection bias when comparing these groups (Schellenberg & Lima, 2024).

Reading enjoyment has been shown to predict reading performance in children (Retelsdorf et al., 2011), likely because greater enjoyment promotes increased reading practice, which in turn strengthens reading skills (Jerrim et al., 2020). In contrast, the current study found no significant relationship between reading enjoyment and lexical decision performance in adults. This discrepancy may reflect developmental differences, as the influence of enjoyment on reading ability may be more pronounced during childhood when literacy skills are still being actively acquired. In adults who have already achieved functional literacy, neural processing mechanisms indexed by cortical tracking may account for variance in reading performance beyond that explained by behavioural factors such as reading enjoyment.

However, we did observe a significant positive relationship between age and reaction times in the lexical decision task, with older participants in our sample responding more slowly. The age range in the current study (19 – 28 years) was relatively narrow, however, this effect could reflect subtle age-related slowing in reaction time even within young adulthood (Baudouin et al., 2004).

### Conclusion

In conclusion, we show that cortical tracking of both speech and music predicts reading performance in a frequency-dependent manner. Stronger delta-band tracking for both speech and music was associated with faster lexical decision reaction times, whereas stronger alpha-band tracking for speech was associated with slower performance, suggesting that optimal reading relies on differential engagement with temporal information across multiple timescales. Importantly, this relationship was not modulated by stimulus type, musical sophistication or reading enjoyment, suggesting that domain-general temporal processing mechanisms, rather than stimulus-specific features, underlie the association between cortical tracking and reading ability. However, causal mechanisms remain unclear, and future research should examine whether training-induced changes in frequency-specific cortical tracking correspond to improvements in reading outcomes.

## Acknowledgments

We thank all participants who so generously gave their time and effort to take part in this study. SCA is funded by the Scottish Graduate School of Social Science (SGSSS) [grant number ES/P000681/1] for her doctoral studies. AK is supported by the Medical Research Council [grant number MR/W02912X/1]. AK is a member of the Scottish-EU Critical Oscillations Network (SCONe), funded by the Royal Society of Edinburgh (RSE Saltire Facilitation Network Award to AK, Reference Number 1963). The funders had no involvement in the study protocol, participant recruitment, data analysis, or manuscript preparation.

## Data availability statement

Data and stimuli are publicly available on the OSF (https://osf.io/xrq36/files/osfstorage).

## Author contributions

S.C.A.: Conceptualisation, methodology, formal analysis, investigation, data curation, writing—original draft, writing—review and editing, visualisation, funding acquisition. S.K.: Investigation, writing – review. S.M.V.: Investigation, writing – review. A.K.: Conceptualisation, methodology, formal analysis, investigation, software, writing – review and editing, supervision, funding acquisition.

## Supplemental material

### Correlation between cortical tracking and word trials in the lexical decision task

To test whether envelope tracking during listening to the story and music predicted participants’ behavioural performance in the lexical decision task, we correlated the MI values per electrode and frequency with participants’ median reaction time across word trials.

For the story condition, we found one negative cluster, indicating that participants who showed stronger envelope tracking to speech at low frequencies had faster reaction times in the lexical decision task (Figure S1A(i), highlighted in red). This negative cluster at low frequencies (0.83 Hz - 1.66 Hz), significantly predicted reaction times in the lexical decision task (Cohen’s *d*_peak_ = 1.79, *p*_cluster_ < .001, 10 electrodes). We also found a positive cluster at higher frequencies (highlighted in black in Figure S1A(i) spanning 10.05 Hz – 13.26 Hz, (Cohen’s *d*_peak_ = 3.11, *p*_cluster_ < .001, 3 electrodes), indicating that participants who showed stronger envelope tracking to speech had slower reaction times in the lexical decision task.

For the music condition, we found one negative cluster, similarly indicating that participants who showed stronger envelope tracking to music at low frequencies had faster reaction times in the lexical decision task (Figure S1A(ii). The negative cluster spanned 0.89 Hz - 1.44 Hz, also predicting reaction times (Cohen’s *d*_peak_ = 2.32, *p*_cluster_ < .001, 12 electrodes). We also found one positive cluster spanning 3.55 – 4.37 Hz (Figure S1A(ii)., indicating that participants who showed stronger envelope tracking to music had slower reaction times in the lexical decision task (Figure S1A Cohen’s *d*_peak_ = 1.24, *p*_cluster_ < .001, 3 electrodes).

### Correlation between cortical tracking and nonword trials in the lexical decision task

To test whether envelope tracking during listening to the story and music predicted participants’ behavioural performance in the lexical decision task, we correlated the MI values per electrode and frequency with participants’ median reaction time across nonword trials.

For the story condition we found three clusters, two negative and one positive. The first negative cluster spanned 1.02 Hz - 1.66 Hz (Figure S1B(i) outlined in black), that predicted reaction times in the lexical decision task (Cohen’s *d*_peak_ = 1.07, *p*_cluster_ < .001, 3 electrodes). The second negative cluster at 7.61 Hz – 8.75 Hz also predicted reaction times in the lexical decision task (Cohen’s *d*_peak_ = 1.30, *p*_cluster_ < .001, 3 electrodes, Figure S1B(i) outlined in yellow), indicating that participants who showed stronger envelope tracking to speech at low frequencies had faster reaction times in the lexical decision task. We also found one positive cluster (Figure S1B(i) outlined in green), indicating that participants who showed stronger envelope tracking to speech at low frequencies had slower reaction times in the lexical decision task. The positive cluster spanned 8.75 Hz – 13.26 Hz (Cohen’s *d*_peak_ = 1.93, *p*_cluster_ < .001, 6 electrodes).

For the music condition, we found one negative cluster in frontal electrodes at low frequencies (0.89 Hz - 1.35 Hz, Figure S1B(ii)) that predicted reaction times in the lexical decision task (Cohen’s *d*_peak_ = 1.84, *p*_cluster_ < .001, 11 electrodes). This indicates that participants who showed stronger envelope tracking to music at low frequencies had faster reaction times in the lexical decision task.

**Figure S1.**
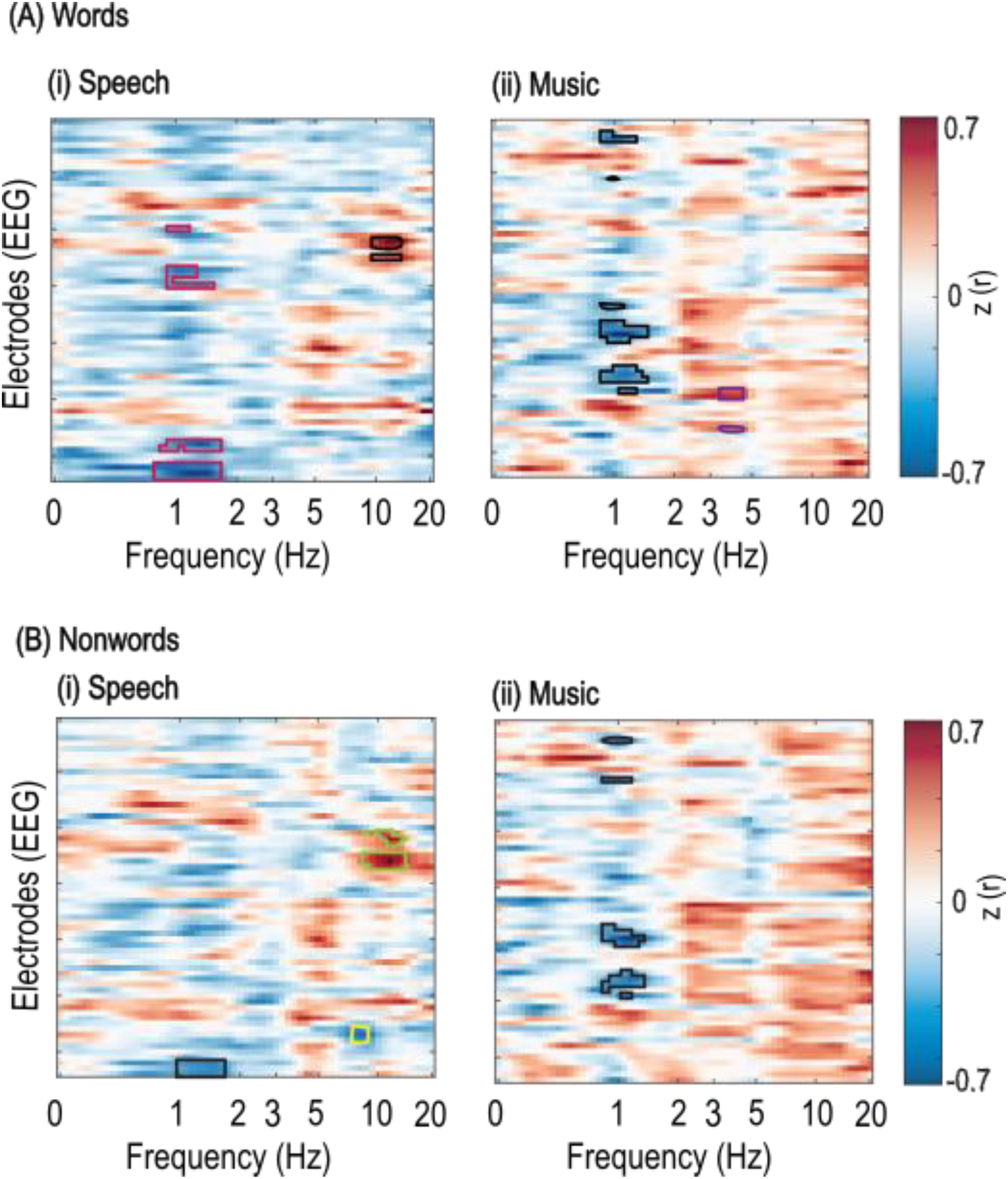
Relationship between cortical tracking and performance on the lexical decision task, quantified through reaction times for word and nonword trials separately. (A) Channel-by-frequency heatmap of the correlation between cortical tracking of speech and participants’ reaction times to word trials. Two clusters were found for speech (i), one negative cluster at low frequencies, and one positive cluster at higher frequencies. Two clusters were found for music (ii): one negative cluster indicating that stronger tracking of music correlated with faster reaction times to words in the lexical decision task, and one positive cluster, indicating that stronger tracking of music at higher frequencies correlated with slower reaction times to words. (B) Channel-by-frequency heatmap of the correlation between cortical tracking of music and participants’ reaction times to nonword trials. Three clusters were found for speech (i), two negative clusters indicating that stronger tracking of speech correlated with faster reaction times to nonwords in the lexical decision task, and one positive cluster at higher frequencies (outlined in green), indicating that stronger tracking of speech correlated with slower reaction times to nonwords. One negative cluster was found for music (ii), indicating that stronger tracking of music correlated with faster reaction times to nonwords in the lexical decision task.

## References

Abrams, D. A., Nicol, T., Zecker, S., & Kraus, N. (2008). Right-hemisphere auditory cortex is dominant for coding syllable patterns in speech. Journal of Neuroscience, 28(15), 3958–3965.

Ahissar, E., Nagarajan, S., Ahissar, M., Protopapas, A., Mahncke, H., & Merzenich, M. M. (2001). Speech comprehension is correlated with temporal response patterns recorded from auditory cortex. Proceedings of the National Academy of Sciences, 98(23), 13367–13372.

Anwyl-Irvine, A. L., Massonnié, J., Flitton, A., Kirkham, N., & Evershed, J. K. (2020). Gorilla in our midst: An online behavioral experiment builder. Behavior research methods, 52(1), 388–407.

Araújo, J., Simons, B. D., Peter, V., Mandke, K., Kalashnikova, M., Macfarlane, A., Gabrielczyk, F., Wilson, A., Di Liberto, G. M., & Burnham, D. (2024). Atypical low-frequency cortical encoding of speech identifies children with developmental dyslexia. Frontiers in Human Neuroscience, 18, 1403677.

Asano, R. (2022). The evolution of hierarchical structure building capacity for language and music: a bottom-up perspective. Primates, 63(5), 417–428.

Atanasova, T., Gross, J., Rimmele, J. M., & Keitel, A. (2026). The involvement of endogenous brain rhythms in speech processing. Neuroscience & Biobehavioral Reviews, 106568.

Baudouin, A., Vanneste, S., & Isingrini, M. (2004). Age-related cognitive slowing: The role of spontaneous tempo and processing speed. Experimental aging research, 30(3), 225–239.

Bhide, A., Power, A., & Goswami, U. (2013). A rhythmic musical intervention for poor readers: A comparison of efficacy with a letter-based intervention. *Mind*, Brain, and Education, 7(2), 113–123.

Boersma, P. (2001). Praat, a system for doing phonetics by computer. Glot. Int., 5(9), 341–345.

Brainard, D. H., & Vision, S. (1997). The psychophysics toolbox. Spatial vision, 10(4), 433-436.

Cason, N., Astésano, C., & Schön, D. (2015). Bridging music and speech rhythm: Rhythmic priming and audio–motor training affect speech perception. Acta psychologica, 155, 43–50.

Chalas, N., Daube, C., Kluger, D. S., Abbasi, O., Nitsch, R., & Gross, J. (2022). Multivariate analysis of speech envelope tracking reveals coupling beyond auditory cortex. NeuroImage, 258, 119395.

De Jong, N. H., & Wempe, T. (2009). Praat script to detect syllable nuclei and measure speech rate automatically. Behavior research methods, 41(2), 385–390.

der Nederlanden, C. M. V. B., Joanisse, M. F., & Grahn, J. A. (2020). Music as a scaffold for listening to speech: Better neural phase-locking to song than speech. NeuroImage, 214, 116767.

Di Liberto, G. M., Peter, V., Kalashnikova, M., Goswami, U., Burnham, D., & Lalor, E. C. (2018). Atypical cortical entrainment to speech in the right hemisphere underpins phonemic deficits in dyslexia. NeuroImage, 175, 70–79.

Ding, N., Patel, A. D., Chen, L., Butler, H., Luo, C., & Poeppel, D. (2017). Temporal modulations in speech and music. Neuroscience & Biobehavioral Reviews, 81, 181–187.

Doelling, K. B., & Poeppel, D. (2015). Cortical entrainment to music and its modulation by expertise. Proceedings of the National Academy of Sciences, 112(45), E6233–E6242.

Drakoulaki, K., Anagnostopoulou, C., Guasti, M. T., Tillmann, B., & Varlokosta, S. (2024). Situating language and music research in a domain-specific versus domain-general framework: A review of theoretical and empirical data. Language and Linguistics Compass, 18(2), e12509.

Flaugnacco, E., Lopez, L., Terribili, C., Zoia, S., Buda, S., Tilli, S., Monasta, L., Montico, M., Sila, A., & Ronfani, L. (2014). Rhythm perception and production predict reading abilities in developmental dyslexia. Frontiers in Human Neuroscience, 8, 392.

Garcia-de-Soria, M. C., Mathias, B., Keitel, A., & Klimovich-Gray, A. (2025). Benefits of music training for learning to read: Evidence from cortical tracking of speech in children. bioRxiv, 2025.2009. 2005.674218.

Giraud, A.-L., & Poeppel, D. (2012). Cortical oscillations and speech processing: emerging computational principles and operations. Nature neuroscience, 15(4), 511–517.

Gorsuch, R. L., & Lehmann, C. S. (2010). Correlation coefficients: Mean bias and confidence interval distortions. Journal of Methods and Measurement in the Social Sciences, 1(2), 52–65.

Goswami, U. (2007). Phonological representations for reading acquisition across languages. In Single-Word Reading (pp. 78-97). Psychology Press.

Goswami, U. (2011). A temporal sampling framework for developmental dyslexia. Trends in cognitive sciences, 15(1), 3–10.

Goswami, U., Huss, M., Mead, N., Fosker, T., & Verney, J. P. (2013). Perception of patterns of musical beat distribution in phonological developmental dyslexia: Significant longitudinal relations with word reading and reading comprehension. Cortex, 49(5), 1363–1376.

Gross, J., Hoogenboom, N., Thut, G., Schyns, P., Panzeri, S., Belin, P., & Garrod, S. (2013). Speech rhythms and multiplexed oscillatory sensory coding in the human brain. PLoS biology, 11(12), e1001752.

Haegens, S., & Golumbic, E. Z. (2018). Rhythmic facilitation of sensory processing: A critical review. Neuroscience & Biobehavioral Reviews, 86, 150–165.

Haegens, S., Händel, B. F., & Jensen, O. (2011). Top-down controlled alpha band activity in somatosensory areas determines behavioral performance in a discrimination task. Journal of neuroscience, 31(14), 5197–5204.

Harding, E. E., Sammler, D., Henry, M. J., Large, E. W., & Kotz, S. A. (2019). Cortical tracking of rhythm in music and speech. NeuroImage, 185, 96–101.

Ince, R. A., Giordano, B. L., Kayser, C., Rousselet, G. A., Gross, J., & Schyns, P. G. (2017). A statistical framework for neuroimaging data analysis based on mutual information estimated via a gaussian copula. Human brain mapping, 38(3), 1541–1573.

Jerrim, J., Lopez-Agudo, L. A., & Marcenaro-Gutierrez, O. D. (2020). Does it matter what children read? New evidence using longitudinal census data from Spain. Oxford Review of Education, 46(5), 515–533.

Keitel, A., Gross, J., & Kayser, C. (2018). Perceptually relevant speech tracking in auditory and motor cortex reflects distinct linguistic features. PLoS biology, 16(3), e2004473.

Keitel, A., Pelofi, C., Guan, X., Watson, E., Wight, L., Allen, S., Mencke, I., Keitel, C., & Rimmele, J. (2025). Cortical and behavioral tracking of rhythm in music: Effects of pitch predictability, enjoyment, and expertise. Annals of the New York Academy of Sciences, 1546(1), 120–135.

Keshavarzi, M., Moore, B. C., & Goswami, U. (2025). Atypical low-frequency and high-frequency neural entrainment to rhythmic audiovisual speech in adults with dyslexia. bioRxiv, 2025.2009. 2014.675562.

Koike, K. J., Hurst, M. K., & Wetmore, S. J. (1994). Correlation between the American Academy of Otolaryngology—Head and Neck Surgery Five-Minute Hearing Test and Standard Audiologic Data. Otolaryngology—Head and Neck Surgery, 111(5), 625–632.

Kotz, S. A., Ravignani, A., & Fitch, W. T. (2018). The evolution of rhythm processing. Trends in cognitive sciences, 22(10), 896–910.

Kumagai, Y., Arvaneh, M., & Tanaka, T. (2017). Familiarity affects entrainment of EEG in music listening. Frontiers in Human Neuroscience, 11, 384.

Lehongre, K., Morillon, B., Giraud, A.-L., & Ramus, F. (2013). Impaired auditory sampling in dyslexia: further evidence from combined fMRI and EEG. Frontiers in Human Neuroscience, 7, 454.

Lehongre, K., Ramus, F., Villiermet, N., Schwartz, D., & Giraud, A.-L. (2011). Altered low-gamma sampling in auditory cortex accounts for the three main facets of dyslexia. Neuron, 72(6), 1080–1090.

Lerdahl, F., & Jackendoff, R. (1983). An overview of hierarchical structure in music. Music Perception, 229-252.

Lubineau, M., Watkins, C. P., Glasel, H., & Dehaene, S. (2024). Examining the Impact of Reading Fluency on Lexical Decision Results in French 6th Graders. Open Mind, 8, 535–557.

Miyagawa, S., Berwick, R. C., & Okanoya, K. (2013). The emergence of hierarchical structure in human language. Frontiers in Psychology, 4, 71.

Molinaro, N., Lizarazu, M., Lallier, M., Bourguignon, M., & Carreiras, M. (2016). Out-of-synchrony speech entrainment in developmental dyslexia. Human brain mapping, 37(8), 2767–2783.

Müllensiefen, D., Gingras, B., Stewart, L., & Musil, J. J. (2013). Goldsmiths Musical Sophistication Index (Gold-MSI) v1. 0: Technical Report and Documentation Revision 0.3. In.

Nora, A., Rinkinen, O., Renvall, H., Arkkila, E., Smolander, S., Laasonen, M., & Salmelin, R. (2024). Impaired cortical tracking of speech in children with developmental language disorder. Journal of neuroscience, 44(22).

Obleser, J., Eisner, F., & Kotz, S. A. (2008). Bilateral speech comprehension reflects differential sensitivity to spectral and temporal features. Journal of Neuroscience, 28(32), 8116–8123.

Obleser, J., & Kayser, C. (2019). Neural entrainment and attentional selection in the listening brain. Trends in cognitive sciences, 23(11), 913–926.

Oldfield, R. C. (1971). The assessment and analysis of handedness: the Edinburgh inventory. Neuropsychologia, 9(1), 97–113.

Olszewska, A. M., Gaca, M., Herman, A. M., Jednoróg, K., & Marchewka, A. (2021). How musical training shapes the adult brain: Predispositions and neuroplasticity. Frontiers in neuroscience, 15, 630829.

Oostenveld, R., Fries, P., Maris, E., & Schoffelen, J.-M. (2011). FieldTrip: open source software for advanced analysis of MEG, EEG, and invasive electrophysiological data. Computational intelligence and neuroscience, 2011.

Osorio, S., & Assaneo, M. F. (2025). Anatomically distinct cortical tracking of music and speech by slow (1–8Hz) and fast (70–120Hz) oscillatory activity. PLoS One, 20(5), e0320519.

Pantaleo, M. M., Arcuri, G., Manfredi, M., & Proverbio, A. M. (2024). Music literacy improves reading skills via bilateral orthographic development. Scientific Reports, 14(1), 3506.

Patel, A. D. (2010). Music, language, and the brain. Oxford university press.

Patel, A. D. (2011). Why would musical training benefit the neural encoding of speech? The OPERA hypothesis. Frontiers in psychology, 2, 142.

Peelle, J. E., & Davis, M. H. (2012). Neural oscillations carry speech rhythm through to comprehension. Frontiers in Psychology, 3, 320.

Pelli, D. G., & Vision, S. (1997). The VideoToolbox software for visual psychophysics: Transforming numbers into movies. Spatial vision, 10, 437–442.

Poeppel, D. (2003). The analysis of speech in different temporal integration windows: cerebral lateralization as ‘asymmetric sampling in time’. Speech communication, 41(1), 245–255.

Power, A. J., Colling, L. J., Mead, N., Barnes, L., & Goswami, U. (2016). Neural encoding of the speech envelope by children with developmental dyslexia. Brain and Language, 160, 1–10.

Retelsdorf, J., Köller, O., & Möller, J. (2011). On the effects of motivation on reading performance growth in secondary school. Learning and instruction, 21(4), 550–559.

Schellenberg, E. G., & Lima, C. F. (2024). Music Training and Nonmusical Abilities. Annu Rev Psychol, 75, 87–128. 10.1146/annurev-psych-032323-051354

Scott, G. G., Keitel, A., Becirspahic, M., Yao, B., & Sereno, S. C. (2019). The Glasgow Norms: Ratings of 5,500 words on nine scales. Behavior research methods, 51(3), 1258–1270.

Smith, Z. M., Delgutte, B., & Oxenham, A. J. (2002). Chimaeric sounds reveal dichotomies in auditory perception. Nature, 416(6876), 87–90.

Sohoglu, E., Peelle, J. E., Carlyon, R. P., & Davis, M. H. (2012). Predictive top-down integration of prior knowledge during speech perception. Journal of Neuroscience, 32(25), 8443–8453.

Swaminathan, S., Schellenberg, E. G., & Venkatesan, K. (2018). Explaining the association between music training and reading in adults. *Journal of Experimental Psychology: Learning*, Memory, and Cognition, 44(6), 992.

Symons, A. E., Dick, F., & Tierney, A. T. (2021). Dimension-selective attention and dimensional salience modulate cortical tracking of acoustic dimensions. NeuroImage, 244, 118544.

Te Rietmolen, N., Mercier, M. R., Trébuchon, A., Morillon, B., & Schön, D. (2024). Speech and music recruit frequency-specific distributed and overlapping cortical networks. Elife, 13, RP94509.

The Mathworks, I. (2025). MATLAB version: 25.1.0.2973910 (R2025a) Update 1. In The MathWorks, Inc. https://www.mathworks.com

Thut, G., Nietzel, A., Brandt, S. A., & Pascual-Leone, A. (2006). α-Band electroencephalographic activity over occipital cortex indexes visuospatial attention bias and predicts visual target detection. Journal of neuroscience, 26(37), 9494–9502.

Tierney, A., & Kraus, N. (2013). Music training for the development of reading skills. Progress in brain research, 207, 209–241.

Wagenmakers, E.-J. (2007). A practical solution to the pervasive problems of p values. Psychonomic bulletin & review, 14(5), 779–804.

Weems, S. A., & Zaidel, E. (2004). The relationship between reading ability and lateralized lexical decision. Brain and Cognition, 55(3), 507–515.

Yao, B., Keitel, A., Bruce, G., Scott, G. G., O’Donnell, P. J., & Sereno, S. C. (2018). Differential emotional processing in concrete and abstract words. Journal of Experimental Psychology: Learning, Memory, and Cognition, 44(7), 1064.

